# Fly wounds and tumors restrict macrophages via a matrix degradation-moderating protease inhibitor

**DOI:** 10.1101/2025.08.12.669954

**Authors:** Kavya Adiga, Yoshiki Sakai, Matin Nawabi, Sofia Mendez-Lopez, Tsai-Ching Hsi, David Bilder

## Abstract

Breaches of epithelial homeostasis trigger an inflammatory response. Not only initiation but negative regulation of the response is critical, as autoinflammation can cause tissue damage and chronic disease. Epithelial breaches can be signaled by damage-associated molecular patterns (DAMPs), including basement membrane (BM) degradation, that attract inflammatory cells. Here we show that the conserved thioester-containing protein Tep3 from Drosophila limits innate immune cell attachment to damaged and transformed epithelia. Tep3 is produced in wounds and tumors alongside the matrix metalloprotease MMP1. Tep3 inhibits MMP1 proteolytic activity, reducing production of a BM DAMP that is necessary for macrophage association. A Drosophila tumor upregulates Tep3 to limit an MMP1- and macrophage-dependent anti-tumor immune response, thus accelerating progression and host death. Hence, fly tumors can exploit a physiological anti-inflammatory axis to pathologically limit their immune restriction.

## INTRODUCTION

Macrophages play central roles in maintaining epithelial homeostasis throughout animal species. Both septic and sterile wounds create signals –pathogen- or damage-associated molecular patterns (PAMPs or DAMPs respectively)--that rapidly recruit macrophages and other innate immune cells, where they mediate the initial defense and repair response. However, this response must be limited to prevent autoinflammatory damage, which is a pathogenic factor in many human diseases^1^. Resolution of inflammation is much less understood than initiation, but involves both the cessation of DAMP and PAMP signals as well as active anti-inflammatory mechanisms. The macrophage response to wounds bears a fascinating but still unclear relationship to the innate immune response to cancer. In mammalian tumor biology, macrophages play a complex role^2,3^. Most studies have examined established tumors, where macrophage infiltration is generally thought to be immunosuppressive by adoption of an M2-like phenotype and by promoting efferocytosis, reducing inflammatory cues. By contrast, there are fewer studies of early tumors where one might expect M1-like macrophage presence to result in increased inflammation and restrain tumor progression. Overall, much remains unknown about such roles in innate tumor immunity, which is a promising frontier for cancer immunotherapy^4,5^.

Drosophila has emerged as an interesting model for studying conserved aspects of macrophage biology and the inflammatory response that involves them ^6,7^. The majority of fly blood cells are called plasmatocytes, which show differentiation via myeloid-like transcription factors, immune-responsive signaling pathways, and phagocytic receptors that are conserved with vertebrate macrophages. These features are sufficiently analogous that plasmatocytes are often referred to as macrophages, as we will do here. Attraction of macrophages to wounds has been extensively studied in the fly embryo, where the main signal involves reactive oxygen species (ROS)^8^. In larvae and adults, the mechanisms are thought to be different: circulating macrophages do not chemotax through the larger hemocoel but instead adhere to wound sites through capture when they are in proximity to DAMP cues^9^. When wound closure is complete, the macrophages disperse, ending the inflammatory response.

Strikingly, macrophages are also attracted to tumors that form in fly larvae. Fly tumors share a number of hallmarks of mammalian cancer, and have become rich systems for studying tumor-host interactions that are conserved between mammals and flies, including lethality-driving syndromes such as cachexia and coagulopathy^10–12^. Macrophages specifically associate with larval fly tumors, with functional consequences that vary depending on the system, including both tumor-promoting and tumor-suppressive outcomes^10,12–24^. Excitingly, these data reveal that invertebrate as well as vertebrate tumors elicit an immune response involving innate inflammatory cells.

While much has been learned about fly inflammation, numerous questions remain about the mechanisms that initiate and limit the cellular immune response, especially beyond the embryo. Here we describe a conserved GPI-linked protease inhibitor that negatively regulates macrophage association with disrupted larval and adult tissues. This molecule, Tep3, is upregulated with and antagonizes a matrix metalloprotease (MMP) that produces a basement membrane (BM)-dependent DAMP, modulating macrophage association with tumors and wounds with functional consequences for animal homeostasis.

## RESULTS

### An adult Drosophila cancer model shows anti-tumor cellular immunity

A cellular immune response to fly tumors has been demonstrated in larval models and transplants, but not in autochtonous adult models. We analyzed a genetically-engineered fly model that creates an ovarian carcinoma (OC) through expression of the oncogenes RasV12 and aPKC^act^ in the follicle cell (ovarian) epithelium, specifically induced in adults via temperature shift^25^. This model has documented grades of tumor progression, from the appearance to initial transformation at grade 1 to severe grade 4 tumors, and displays several tumor-host interactions conserved with human patients. We analyzed macrophages using an antibody against NimC1, a phagocytic receptor found on most of these cells. In wild-type (WT) ovaries, only a few macrophages are found; these are located at the germarium and display a compact, rounded morphology (**Fig. 1A, A’**); macrophages are also seen on the posterior oviduct. By contrast, tumorous ovaries show a dramatic increase in macrophages, which are primarily associated with the transformed cells at the anterior ovary (**Fig. 1B, C**). Tumor-associated macrophages (TAMs) have a spread morphology with multiple cellular extensions, a phenotype previously termed ‘macrophage activation’ (**Fig. 1B’**)^26^.

**Figure 1:**
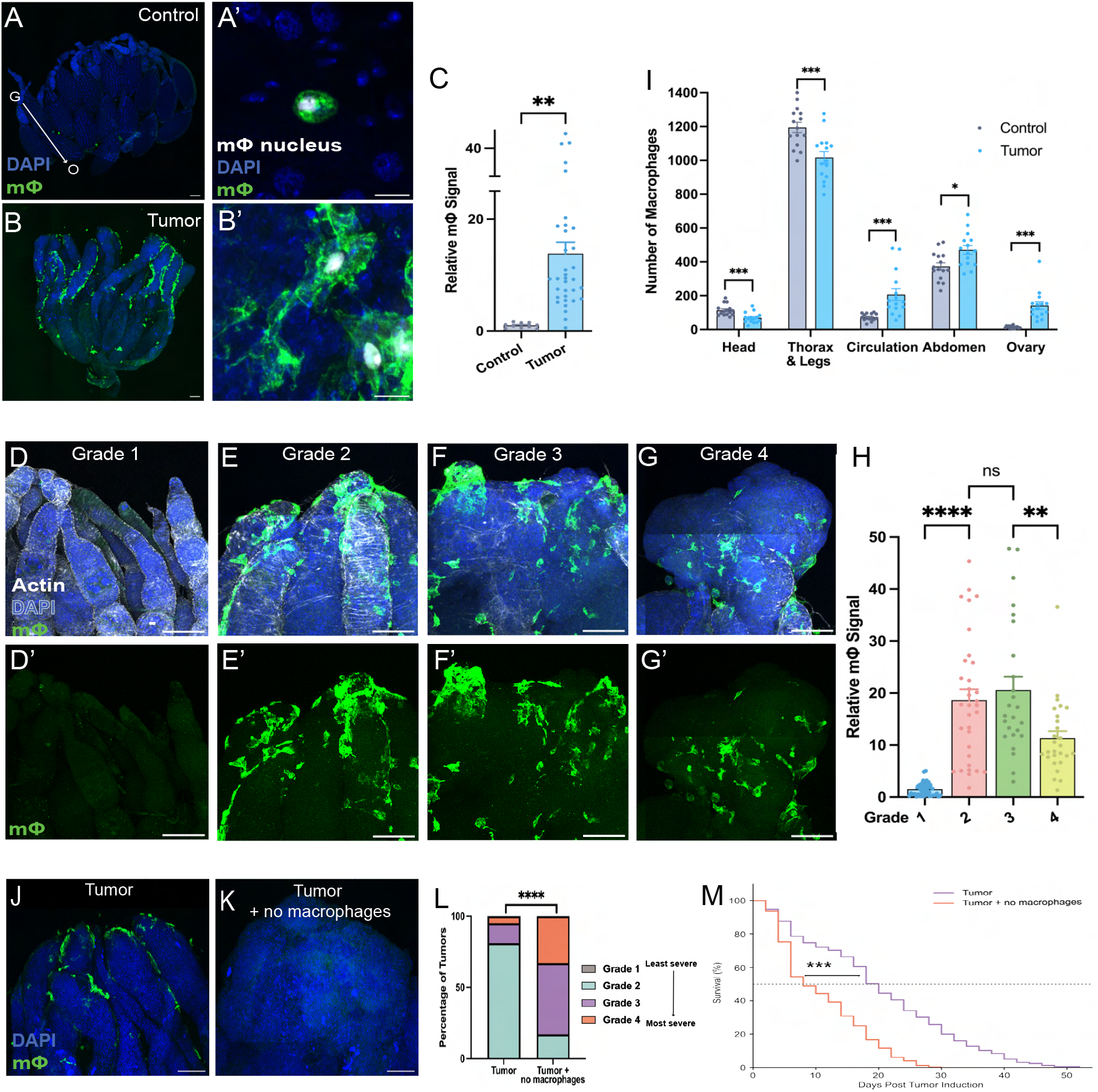
Adult macrophages recognize tumors and slow their progression. (**A, B)** OC tumors exhibit increased macrophages detected by anti-NimC1 (green), quantitated in **C** (n-values: Control = 10 Tumor = 34). Control macrophages are round (**A’**) while TAMs display a spread morphology associated with macrophage activation (**B’**). (**D-G**) TAMs are not found on Grade 1 tumors but start to bind at Grade 2 and continue through Grades 3 and 4, quantitated in **H** (n-values: Grade 1 = 45, Grade 2 = 33, Grade 3 = 25, Grade 4 = 27). (**I**) Tumor-bearing flies at day 10 ATI have fewer thorax and head sessile macrophages with an increase in abdominal, circulating, and tumor-associated macrophages. (**J, K**) Tumor-bearing flies with no macrophages show faster tumor progression and accelerated host death compared to tumor-bearing flies with macrophages, quantitated in **L, M** (n-values (L): Tumor = 58, Tumor + no macrophages = 24; (M): Tumor = 155, Tumor + no macrophages = 239). Scale bars = 100μm (A, B, D-G, J, K), 10μm (A’, B’); Unpaired t-test (C, I), one-way ANOVA (H), Mann-Whitney U-test (L), and log-rank test (M) used to determine significance; *p < .05, **p< .005, ***p < .0005, ****p < .00005. Dots in (C), (H), (I) represent individual flies.

We examined the kinetics of TAMs with respect to OC progression (**Fig. 1D-H**). TAMs were first seen on grade 2 tumors, as evidenced by a 20-fold increase in NimC1 reactivity. Elevated macrophage numbers continued in grade 3 and grade 4 tumors. We examined circulating and sessile macrophage populations and found that the increase in TAMs correlated with a decrease in the sessile population from the thorax and head but not the abdomen (**Fig. 1I**), indicating that a specific macrophage subpopulation is mobilized for recruitment to the tumor.

To test a functional role in the OC model, we generated flies in which macrophages were ablated through GAL4-driven expression of apoptosis inducers specifically in adults. OC tumors were driven by LexA-based expression of RasV12 and aPKC^act^ in follicle cells, which produces tumors similar to those driven by GAL4. The ablation system used eliminates most macrophages^27^, and indeed no TAMs were seen in these flies (**Fig. 1K**). Macrophage-depleted flies showed strongly accelerated tumor progression, with a ∼6.5-fold increase in grade 4 tumors and a nearly 4-fold increase in grade 3 tumors at day 15 after tumor induction (ATI), compared to macrophage-replete control flies (**Fig. 1J-L**). Macrophage-depletion also accelerated the death of tumor-bearing flies (**Fig. 1M**). Therefore the OC model induces a cellular immune response in the adult that has a net anti-tumor activity.

### Ovarian tumors express a protein that restricts tumor-associated macrophages

To explore mechanisms that could regulate macrophages in OC flies, we tested oncokines: secreted factors that are upregulated in the tumor and could mediate communication with non-tumor cells. Amongst a list of oncokines from RNA sequencing of OC tissue, including those previously validated to mediate tumor-host interactions^10,25,28,29^, we focused on Thioester-containing protein 3 (Tep3). *Tep3* is upregulated ∼12-fold in the OC transcriptome (**Fig. 2D**)^25^. We confirmed upregulation by examining a *Tep3-GAL4* transcriptional reporter in OC tumors. *Tep3* expression was not evident in control ovaries but was first elevated in subsets of transformed cells within grade 1 tumors, with expression increased in tumors of grades 2 and higher (**Fig. 2A-C**).

**Figure 2:**
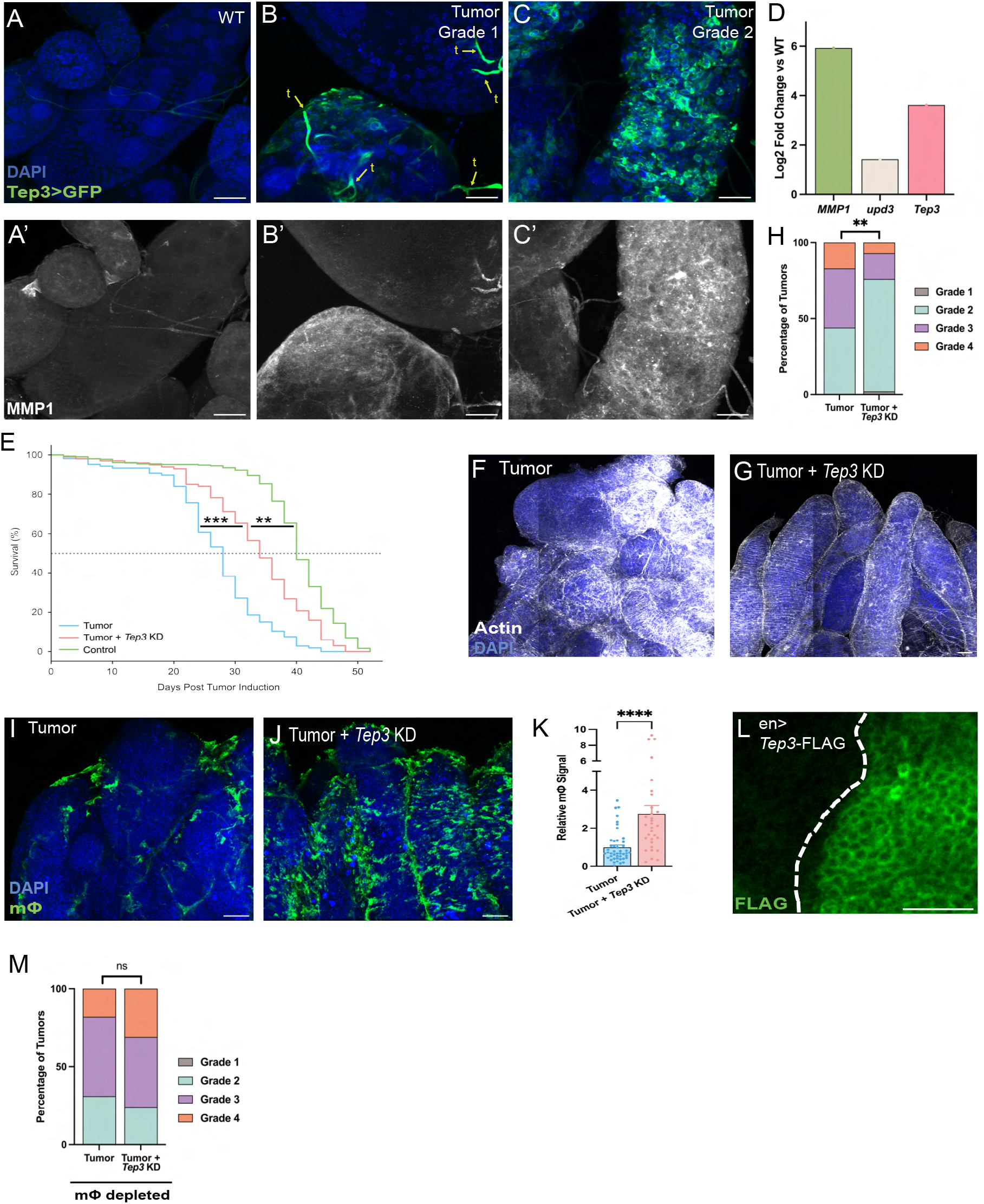
Tumor-produced Tep3 limits macrophage association. (**A-C**) *Tep3* levels are increased in tumors compared to control ovaries (green). *Tep3* is also expressed in WT trachea (t). (**A’-C’**) *Tep3* expression is correlated with MMP1 antibody staining. (**D**) OC tumors upregulate *Tep3* alongside expected genes such as pro-invasion *MMP1* and pro-inflammation *upd3*. Tumors with *Tep3* knock down (KD) display increased host lifespan (**E**) and slower tumor progression (**F, G**) (DAPI (blue) and actin (white)), quantitated in **H** (n-values: (E) Tumor = 107, Tumor + *Tep3* KD = 87; (H) Tumor = 36, Tumor + *Tep3* KD = 53). (**I, J**) Tumors with *Tep3* KD exhibit increased TAMs (green), quantitated in **K** (n-values: Tumor = 45, Tumor + *Tep3 KD* = 31). (**L**) Overexpression of FLAG-tagged Tep3 in wing disc portion (marked by dotted white line) shows membrane expression (green). (**M**) Macrophage-depleted flies show no significant difference in tumor progression between WT and *T*e*p3 KD* tumors, indicating that Tep3 effect on tumor progression is macrophage-dependent. (n-values: Tumor = 39, Tumor + *Tep3* KD = 71). Scale bars = 50μm (F, G, I, J), 20μm (A, A’, B, B’, C, C’, L); Unpaired t-test (K), Mann-Whitney U-test (H, M), log-rank test (E) used to determine significance; **p < .005, ***p < .0005, ****p < .00005. Dots in (K) represent individual samples.

For functional tests, we depleted Tep3 via RNAi specifically in the follicle cells alongside the two oncogenes. Importantly, Tep3 depletion in tumors induced a strong extension of the lifespan of tumor-bearing flies (**Fig. 2E**). Analysis of tumor grade revealed that Tep3-depleted tumors progressed more slowly that those depleted for a control RNAi construct, consistent with the lifespan extension (**Fig. 2F-H**). Extension was also seen with a second independent RNAi construct (**Fig. S1A**). Thus, tumor-produced Tep3 accelerates tumor progression and host death.

Since depleting either macrophages or Tep3 leads to opposite effects on tumor progression and lifespan, we next analyzed TAMs. Strikingly, Tep3-depleted tumors displayed nearly 3-fold more TAMs than control tumors (**Fig. 2I-K**). The difference was independent of tumor grade: that is, both grade 2 and grade 3 Tep3-depleted tumors contained more TAMs than control counterparts. Tep3 depletion had no impact on macrophage numbers on otherwise WT ovaries (**Fig. S1D**). This suggested that Tep3 is produced by the tumor to limit macrophage association. We tested the link between Tep3-regulated macrophage association and Tep3-regulated tumor progression by depleting Tep3 in OC tumors while eliminating macrophages with apoptotic inducers using LexA-driven expression. When macrophages were killed, Tep3 depletion lost its ability to slow tumor progression; there was no significant difference in tumor grades between Tep3-expressing and Tep3-depleted tumors (**Fig. 2M**). These results suggest that Tep3 acts via macrophages to limit the host anti-tumor immune response.

### Basement membrane digestion cues macrophages to attach to the tumor

How could Tep3 negatively regulate macrophage association with OC tumors to accelerate progression and host death? TEP family members are found across both vertebrate and invertebrate phyla; perhaps the best known are vertebrate alpha-2-macroglobulin (a2M) proteins that are plasmatic protease inhibitors and function in inflammation and immunity^30^. Drosophila contains 4 secreted TEP family members that are related to a2M. Insect TEPs have been previously investigated for immune roles, including action as opsonins or participating in complement pathways^31^. Tep3 is perhaps the least studied, and may not be induced by microbial challenge like Teps 1, 2 and 4^32–35^. Tep3 is also unique amongst fly TEPs in having a predicted GPI anchor, suggesting a local rather than systemic role. Interestingly, the predicted mammalian ortholog of Tep3 is CD109 (**Fig. S1B-C**), which, unlike a2Ms or the related complement C3/C4-like proteins, also has a GPI anchor^36^. Expression of a tagged transgene confirmed Tep3 membrane enrichment (**Fig. 2L)**.

A familiar paradigm for macrophage recognition of wounded or transformed tissue involves DAMPs that signal the presence of dying or unwanted cells, triggering their removal^37,38^. ROS is one characterized DAMP to which both fly and vertebrate macrophages respond, while dying cells can release a variety of components that have DAMP activity. We found no difference in apoptosis between OC tumors with and without Tep3 depletion at day 10 ATI (**Fig. 3A-C**). Inducing cell death in non-tumorous ovaries was not sufficient to attract macrophages, and blocking cell death in OC tumors through overexpression of DIAP1 or p35 also had no effect on TAM numbers (**Fig. S2D-H**). The same was true for OC tumors in which ROS was reduced by depletion of Dual Oxidase (**Fig. S2I-L**).

**Figure 3:**
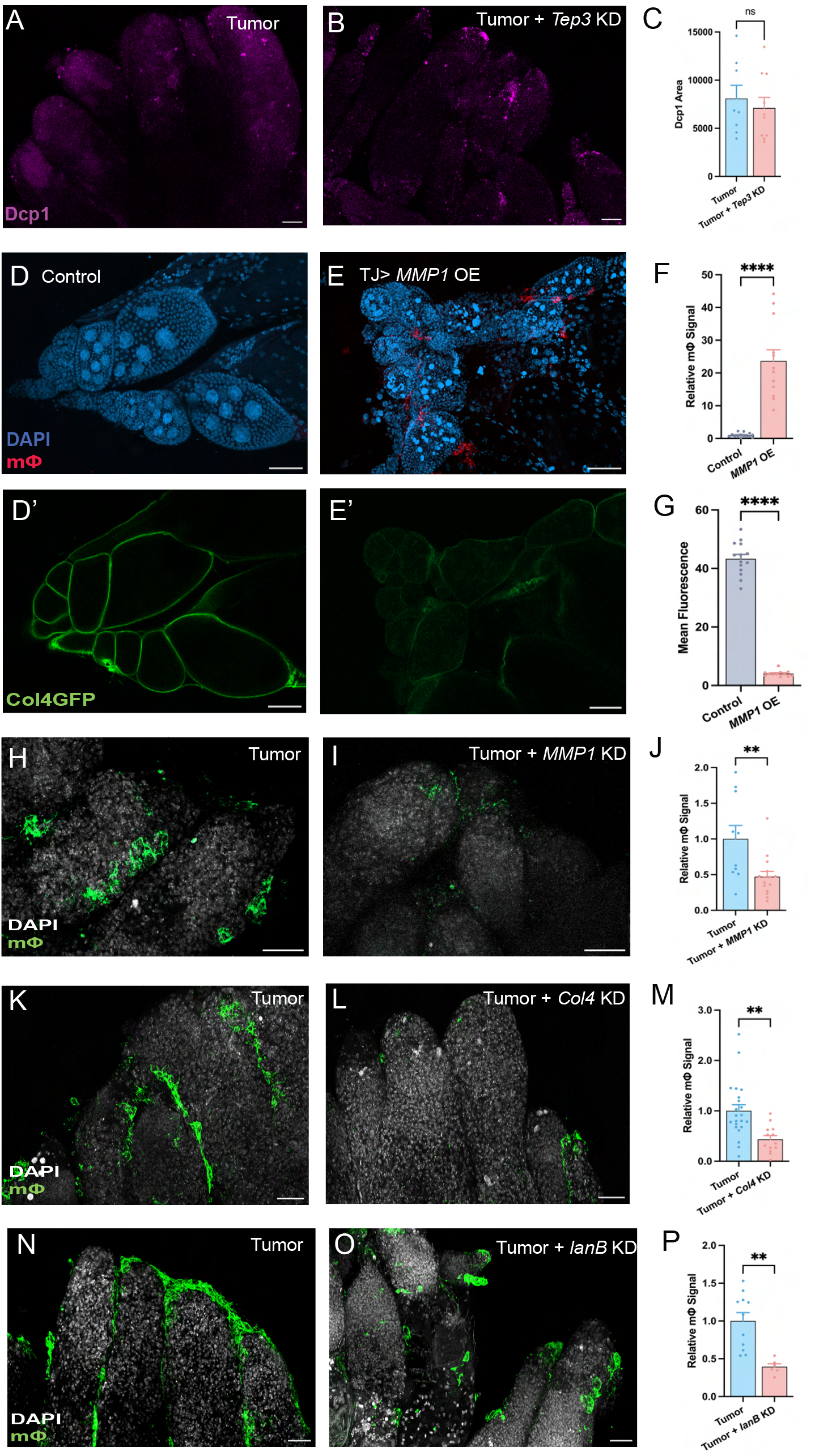
Macrophage association with tumor involves basement membrane breakdown. (**A-B**) *Tep3* KD in tumors does not affect apoptosis at day 10 ATI, indicated by anti-Dcp1 staining (magenta), quantitated in **C** (n values: Tumor = 29, Tumor + *Tep3* KD = 16). (**D-E**) *MMP1* overexpression in a WT ovary is sufficient to recruit macrophages (red) and reduce BM levels (**D’-E’**, green, single confocal slice), quantitated in **F-G** (n-values: (F) Control = 14, *MMP1* OE = 12; (G) Control = 14, *MMP1* OE = 12). **(H-I)** Tumors with *MMP1* KD exhibit decreased TAMs (tumor marked by dotted white line, macrophages in green), quantitated in **J** (n-values: Tumor = 10, Tumor + *MMP1* KD = 15). Depleting Col4 **(K-L)** or Laminin **(N-O)** in OC tumors reduced TAMs (green), quantitated in **(M, P)** (n-values: (M) Tumor = 22, Tumor + *Col4* KD = 13; (P) Tumor = 11, Tumor + *lanB* KD = 6). (Scale bars = 50μm; Unpaired t-test (C, F, G, J, M, P) used to determine significance; **p < .005, ****p < .00005. Dots in (C), (F), (G), (J), (M), (P) represent individual samples.

In fly larvae, breaches of the basement membrane (BM) due to either mechanical disruption or enzymatic degradation result in macrophage attachment^14,15,18,39^. To our knowledge past experiments do not distinguish whether macrophages recognize the absence of BM components on tissue per se, or alternatively the activity of BM-degrading enzymes such as MMPs. Since follicle cells produce most of their own BM^40^, we first depleted the BM components Collagen IV (Col4) or Laminin from otherwise WT ovaries and found that depletion of Col4 or Laminin, caused a minor (2-fold) increase in macrophages, predominantly in the ‘companion’ population at the germarium (**Fig. S3A-F**). We then induced BM degradation through transgenic expression of either of the two MMPs encoded in the fly genome, named MMP1 and MMP2. While ovaries overexpressing MMP2 could not be recovered, overexpression of MMP1 in otherwise WT follicle cells was sufficient to reduce BM levels and cause a strong 20-fold increase in macrophage attachment (**Fig. 3D-G**).

We next analyzed the OC model. Previous transcriptome data showed that *MMP1* is upregulated in these tumors (**Fig. 2D**)^25^, and BM levels are reduced (**Fig. 4A-D**)^25^. To functionally test a role for MMP1, we depleted it from OC tissue. At day 10 ATI, when TAMs are first seen on control tumors, MMP1-depleted tumors showed decreased TAM numbers, with no significant change in tumor grade compared to control tumors (**Fig. 3H-J**). Importantly, when Col4 or Laminin was depleted in OC, decreased TAMs were also seen (**Fig. 3K-P**). Together, these data indicate that tumor MMP1 digests BM to release the relevant epithelial DAMP for macrophage association.

**Figure 4:**
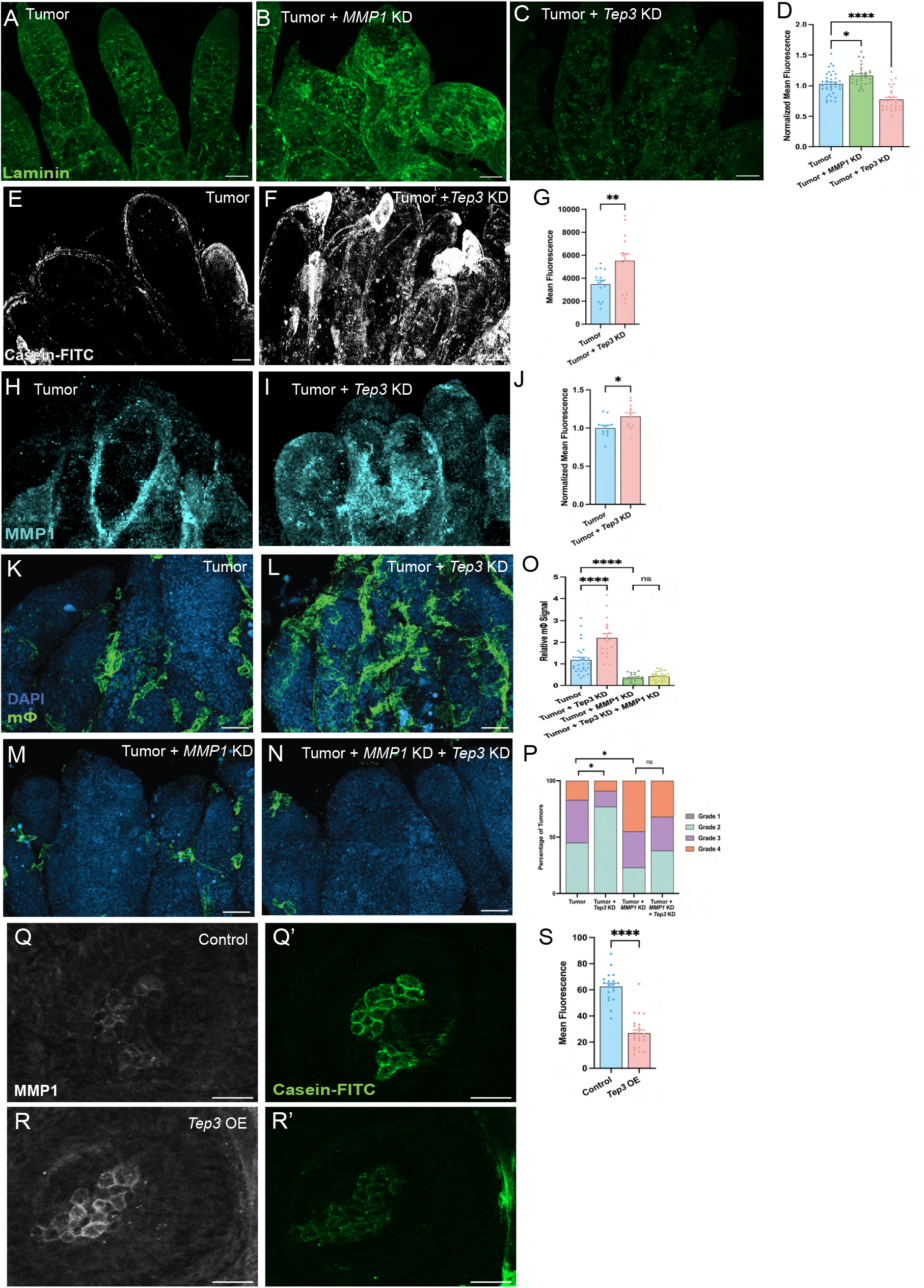
Tep3 inhibits MMP1 activity to modulate immune recognition of tumor. Compared to control tumors (**A**), tumors with *MMP1* KD show increased BM (anti-Laminin antibody, green) (**B**), while tumors with *Tep3* KD display decreased BM (**C**), levels quantitated in **D** (n values: Tumor = 37, Tumor + *MMP1* KD = 32, Tumor + *Tep3* KD = 30). (**E-F**) Tumors with *Tep3* KD display increased MMP1 activity, indicated by Casein-FITC fluorescence (white), quantitated in **G** (n values: Tumor = 14, Tumor + *Tep3* KD = 15). (**H-I**) *Tep3* KD tumors have slightly increased MMP1 levels, (cyan), quantitated in **J** (n values: Tumor = 12, Tumor + *Tep3* KD = 12). (**K-N**) Tep3 function in tumors depends on MMP1, as TAM increase (green) in *Tep3 KD* tumors compared to control tumors (**K, L**) is lost with *MMP1* and *Tep3* double KD, which display TAM numbers similar to tumors with *MMP1* KD alone (**M, N**), quantitated in **O** (n-values: Tumor = 29, Tumor + *Tep3* KD = 19, Tumor + *MMP1* KD = 20, Tumor + *Tep3* KD + *MMP1* KD = 28). This dependence is also reflected in tumor progression where the slower tumor progression caused by *Tep3* KD is lost upon combined KD of *Tep3* and *MMP1* **(P)** (n-values: Tumor = 40, Tumor + *Tep3* KD = 44, Tumor + *MMP1* KD = 44, Tumor + *Tep3* KD + *MMP1* KD = 53). **(Q-S)** WT larval eye disc macrophages express MMP1 (white) and display Casein cleavage activity (green); Tep3 overexpression in macrophages reduces MMP1 activity (quantitated in **S**) but not protein levels. (n values: Control = 20, *Tep3* OE = 24). Scale bars = 50μm (A, B, C, E, F, H, I, K, L, M, N), 20μm (Q, Q’, R, R’); Unpaired t-test (G, J, S), Mann-Whitney U-test (P), and One Way ANOVA (D, O) used to determine significance; *p < .05, **p < .005, ****p < .00005. Dots in (D, G, J, O, S) represent individual samples.

### Tep3 inhibits MMP1 activity to modulate macrophage association

The activity of TEP family proteins as protease inhibitors suggested a hypothesis that Tep3 inhibits MMP1 activity to regulate TAMs in the OC model. Immunostaining for MMP1 protein showed elevation first in some cells of grade 1 tumors (**Fig. 2B**). Interestingly, these MMP1-positive cells often coexpressed the *Tep3* transcriptional reporter. At later stages, MMP1 became more widespread throughout the tumor, as did *Tep3* (**Fig. 2C-D**). Strikingly, BM levels were significantly lower in Tep3-depleted compared to control grade-matched tumors (**Fig. 4A-D**), suggesting the hypothesis that Tep3 might oppose MMP1-mediated BM destruction.

Since TEP proteins inhibit protease activity rather than protease levels, we turned to a fluorometric assay. MMP1 protease activity can be detected by liberation of FITC otherwise quenched by casein, a demonstrated MMP1 substrate^41^. When incubated with casein-FITC, tumors revealed FITC dequenching while WT ovaries did not, correlating with MMP1 immunostaining (**Fig. S3G-I**). We then employed the casein-FITC assay on Tep3-depleted tumors. Fluorescence quantitation showed a clear increase in grade-matched tumors compared to tumors depleted with a control RNAi (**Fig. 4E-G**). We examined whether lower MMP1 enzymatic activity was due to MMP1 protein downregulation and found a slight but significant increase in Tep3-depleted vs control tumors (**Fig. 4H-J**). We also tested whether Tep3-depleted tumors altered gelatin-FITC fluorescence in an analogous assay validated for MMP2 activity^41^. No difference was seen (**Fig. S4A-C**).

An implication of the hypothesis that tumor-produced Tep3 restricts MMP1 activity is that MMP1 produces the DAMP sensed by OC TAMs. We therefore examined OC tumors that were depleted simultaneously for both MMP1 and Tep3. These tumors showed no significant difference in TAMs compared to MMP1 depletion alone (**Fig. 4K-O**). By day 20 ATI, MMP1-depleted tumors showed faster tumor progression than control tumors, and the same accelerated progression was seen in MMP1 Tep3 double-depleted tumors (**Fig. 4P**). Thus, upregulated MMP1 increases TAM numbers, and Tep3 requires MMP1 to limit this phenotype.

To further test the relationship between Tep3 and MMP1, we sought an independent site of native, non-pathological MMP1 expression. We noticed that macrophages associated with the eye imaginal disc of WT L3 larvae expressed MMP1 (**Fig. 4Q**) and, as expected, consistently liberated casein-FITC (**Fig. 4Q’**). However, driving transgenic expression of Tep3 in hemocytes reduced casein cleavage in the fluorometric assay, with no reduction in MMP1 protein levels (**Fig. 4R-S**).

### Wound-induced Tep3 inhibits MMP1 in physiological contexts

Why would a tumor produce both a BM-cleaving MMP and an inhibitor of that enzyme? Since tumors often activate physiological pathways in pathological contexts, we considered what the normal functions of Tep3 might include. Epithelial wounds share many characteristics with tumors, including disruption of tissue structure and attachment of macrophages^42,43^. We therefore wounded imaginal discs by physically pinching them, both ex vivo and in vivo. In ex vivo culture, MMP1 was upregulated at the wound site ∼3 hours after damage, alongside pyknotic nuclei typical of dying cells (**Fig. 5B-D**). Remarkably, MMP1 colocalized here with new induction of the *Tep3* transcriptional reporter, which, as in unwounded discs, also showed expression in the future wing blade (**Fig. 5A-B**). The FITC-casein assay revealed MMP1 activity along the wound site (**Fig. 5B-D**). When Tep3 was overexpressed in the anterior compartment of the disc, it caused a significant reduction of this activity, with no reduction in MMP1 levels (**Fig. 5E-H)**. Moreover, fewer macrophages were recruited to the wound *in vivo* (**Fig. 5I, J**). Thus, Tep3 regulation of MMP1 and macrophage attachment in genetically altered tumors reflects a normal action in preserving homeostasis of WT animals.

**Figure 5:**
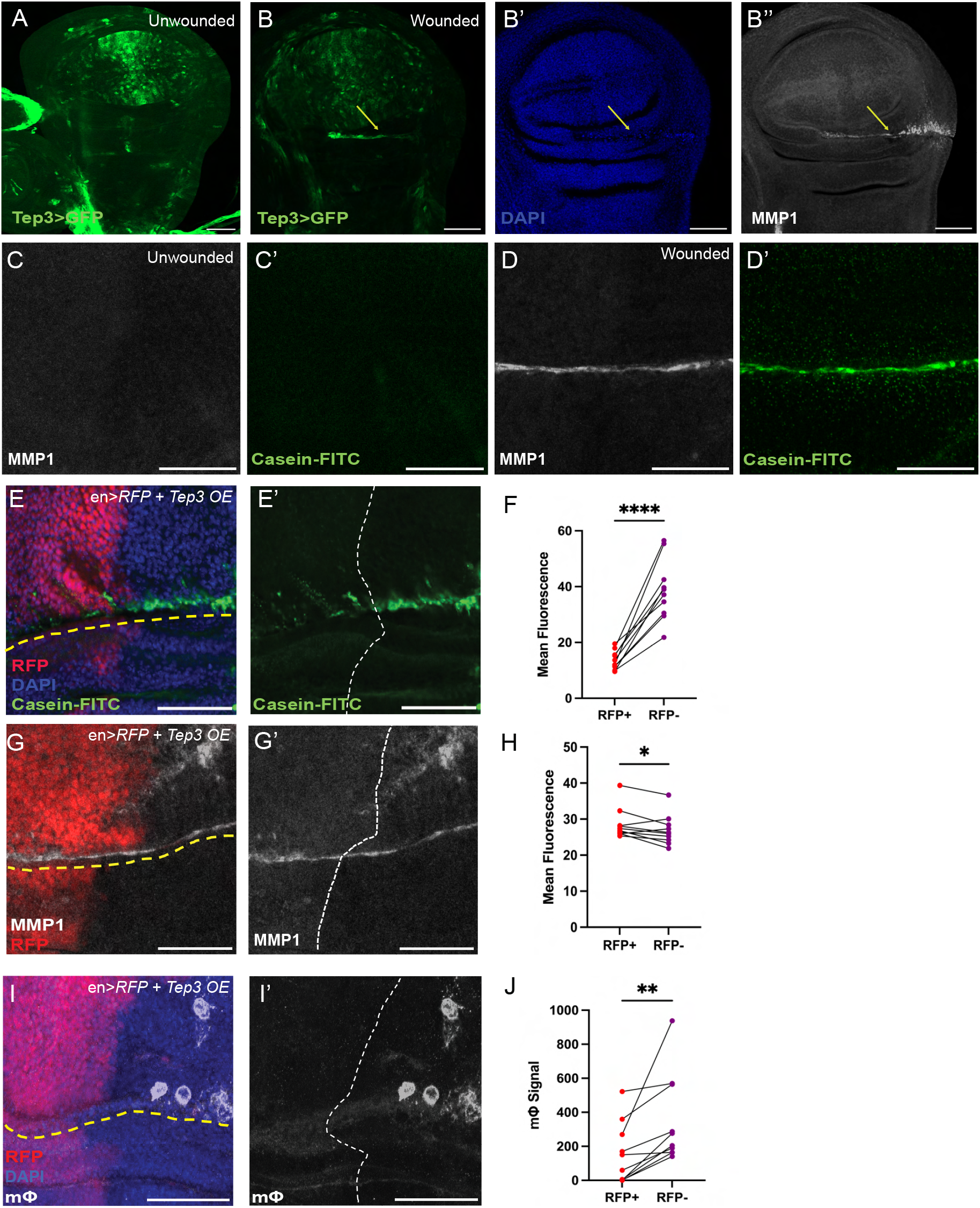
Tep3 antagonizes MMP1 in wounds. **(A)** WT larval wing discs express *Tep3* (*Tep3 GAL4>GFP*, green) in the future wing blade. **(B)** Wounded wing discs (marked by yellow arrow, with pyknotic nuclei revealed by DAPI in blue **(B’)**) show induction of MMP1 (white) **(B’’)**, and colocalized *Tep3* (green). **(C-D)** Wound MMP1 (white) is correlated with increased Casein cleavage (green), indicating elevated MMP1 activity. Overexpressing *Tep3* in the anterior compartment of a wounded wing disc (*engrailed*-expressing *en+* region in red, wound marked by yellow line) specifically reduces MMP1 activity (green) **(E-E’)** in *Tep3-*expressing wound cells without reducing MMP1 levels (white) **(G-G’**), quantitated in **F, H** (n values: (F) 10, (H) 10). Macrophage association (white) **(I-I’)** is also decreased in the *Tep3-*expressing region, quantitated in **J** (n values: 10). Scale bars = 20μm; paired t-test used to determine significance; *p < .05, **p<.005, ****p < .00005. Dots in (F, H, J) represent individual samples.

## DISCUSSION

Mechanisms that regulate immune cell recruitment are central to both physiological and pathological inflammatory responses. We show here that fly Tep3 is a macrophage association-limiting molecule that is induced upon epithelial disruption. TEPs from many species have immune- or inflammation-related roles, including the mammalian proteins C3/C4 that function in the complement system and alpha-2-macroglobulin which is released during tissue injury^30^. Previously proposed functions for insect TEPs include action as opsonins or participation in a complement-related pathway^35,44–46^. While these remain possibilities for the infection-induced secreted Drosophila Tep1, Tep2 and Tep 4, they are unlikely to be the case for GPI-linked Tep3, which seems to have a local mechanism of action. Prior work on Drosophila TEPs suggested that *Tep3* mutants are viable and have limited phenotypes, including no sensitivity to wounding^33,34^. However, the allele studied does not delete coding sequence, and we find that it acts as a hypomorph and that *Tep3* is in fact an essential gene (**Fig. S4D**). Conclusions about Tep3 function derived from this allele should be revisited.

Tep3 is upregulated in both tumors and wounded tissue, and works at these sites to antagonize MMP1, as demonstrated by the MMP1-dependence of examined *Tep3* phenotypes. Like inflammatory MMP1, anti-inflammatory Tep3 induction may be activated by the wound-induced JNK pathway, and indeed the two proteins are coexpressed in a number of contexts; inspection of datasets reveals JNK-dependent *Tep3* upregulation alongside *MMP1* in larval tumors as well as regenerating wounds^47,48^. We suggest that Tep3 expression is normally dynamically tuned to limit MMP1 activity to appropriate levels, preventing excess BM degradation and helping to resolve inflammation.

A direct target of Tep3 inhibitory activity remains unknown. Data are consistent with, but do not demonstrate, that MMP1 is a substrate. Some TEPs inhibit targets through forming covalent bonds, while other TEPs use a non-covalent mechanism for protease inhibition, which hampers biochemical identification of substrates^30,49^. A related unanswered question is the identity of the DAMP processed by MMP1 to induce macrophage attachment. One possibility, suggested by the requirement for Col4 and Laminin in this MMP1-dependent phenotype, is that fly macrophages detect digested BM fragments. Mammalian macrophages can sense such fragments as DAMPs through receptors such as RAGE and CD44 that have no obvious homolog in flies, as well as integrins, CD36, and TLR2/4 which do have fly homologs^37,38,50^. The identification of Tep3 and MMP1 substrates in this inflammatory axis is an important future goal.

This work highlights a number of similarities between Tep3 and its mammalian ortholog, CD109, which is also a uniquely GPI-linked TEP (reviewed in ^36,51^). CD109 is upregulated in tissue damage, and is overexpressed in various carcinomas as well as glioblastoma; high CD109 levels indicate poor prognosis in multiple cancers. Experimental overexpression of CD109 decreases inflammation, while CD109 mutant mice show excess immune cell infiltration. CD109 phenotypes also point to an ECM-regulating role: mutant mice exhibit fibrosis, while overexpression increases collagen organization. These effects have been largely attributed to inhibition of TGFB signaling through direct interaction with both receptors and ligands, but this is not the case in all cell types. The fly results suggest that MMP antagonism is another CD109 role that merits examination.

Our data extend the cellular immune response to tumors to genetically engineered adult flies, and demonstrate that in the OC model macrophages have an anti-tumor activity. Ablation of macrophages in these animals accelerates progression, while a condition that recruits more macrophages to the tumor slows progression. Previous work in larval and allograft models has revealed varying roles for macrophages, from pro-tumor to anti-tumor to not involved^10,12,13,16–24^. Suggested mechanisms of anti-tumor macrophage activity include death caused by the TNF Eiger, ROS, or efferocytosis of transformed cells. However, we do not see enhanced cell death in high-grade OC tumors depleted for Tep3 (**Fig. S2A-C**), and death is not restricted to TAM-proximal sites. OC progression involves in part increased tumor invasiveness, and one role of macrophages in wound repair is to secrete BM components. Thus a repair-like contribution of BM components is one potential mechanism by which TAMs could limit tumor progression.

Both similarities and differences between human and fly tumors are revealed by this study. In both organisms, infiltration by immune cells is a positive indicator of an effective anti-tumor response, and such infiltration can negatively correlate with ECM levels in the tumor microenvironment (TME). However, in humans, macrophages are often present in the TME of ‘cold’ tumors that exclude tumorlytic T cells^2,3^. In the fly, where macrophages are the sole immune cells, OC TAM numbers correlate with the strength of the anti-tumor response and host prognosis. Additionally, the best-known DAMPs produced by human wounds and tumors are cytosolic molecules released by necrotic death, stimulating an inflammatory reaction, whereas apoptosis is generally considered immunosuppressive^37^. Blocking apoptosis did not influence TAM presence in the OC model, and OC tumors show limited necrosis; there is little information about cytosolic DAMPs in flies. Notably, the DAMP implicated by our work --BM damage through JNK-regulated MMP production--is agnostic to the type of cell death caused by a lesion, but is one of many common features shared by both tumors and wounds^42,43^. In wounds, JNK signaling resolves as tissue repair is completed, but in fly tumors JNK is constitutively driven by oncogenic activity. In the OC model, this leads to constant high levels of Tep3, which blunts the anti-tumor immune response. Thus, tumors can exploit a physiological axis that reduces inflammatory cell association to promote their progression and kill the host more rapidly.

## Acknowledgements

We thank I. Ando, A. Herzig, K. Bruckner, B. Lemaitre, J. Shim and D. Montell for generous gifts of reagents, as well as the community resources provided by the Bloomington Drosophila Stock Center (NIH P40OD018537), TRiP at Harvard Medical School (NIH/NIGMS R01-GM084947), and the Kyoto stock center. We are grateful to L. Mathies for generating *UAS-Tep3-FLAG*, S. Haraguchi for initial experiments characterizing *Tep3*, and the entire Bilder lab for helpful discussions. This work was supported by a Mark Foundation ASPIRE Award, American Cancer Society grant DBG-24-1322649, and NIH grant GM130388 to D.B., as well as Cancer Research Coordinating Committee Fellowships to K.A and T.H. and a Damon Runyon Cancer Research Foundation Fellowship to Y.S.

## EXPERIMENTAL MODEL AND STUDY PARTICIPANT DETAILS

### Fly husbandry and genotypes

Flies were maintained at 21°C in wide vials on cornmeal, molasses, and yeast food. *UAS-Tep3-FLAG* places 3X FLAG tags C-terminal to the *Tep3* coding sequence in the *pUASTattB* vector and was inserted into the *attP2* site via microinjection (BestGene Inc). For targeting *Tep3*, RNAi construct #63660 from BDSC was used except in Fig. S1A. In RNAi or overexpression experiments, controls were utilized with matched *attP* sites targeting GFP or another exogenous protein. Complete genotypes are provided in **Table S1**. For *Tep3* mutant alleles, *Tep3 sk2* is a 23 bp deletion that creates a frameshift in the fifth exon, terminating after the first quarter of the protein; *Tep3-GAL4* is a CRIMIC inserted in the first intron which is expected to terminate the protein shortly after initiation; *Tep qD* deletes ∼1.5 kb of intragenic sequence ∼250 bp 5’ to the *Tep3* start codon^33^.

For the LexA-driven OC model, the sequence encoding the chimeric transcriptional activator LexA-Hinge-GAL4 Activation Domain (LHG) was inserted using CRISPR/Cas9 HDR at the *tj* locus, in the same site as the widely used *tj-GAL4* insertion. This chromosome was then recombined with *tubGAL80ts. LexAop-aPKCact + RasV12* was created by cloning sequences encoding aPKCdelN and Ras85DV12, separated by the T2A self-cleaving peptide, into *pJFRC19*. The transgene was inserted into the *attP2* site. To build *Hml*^*P2A*^*-LHG*, the *Hml*^*P2A*^ promoter sequence from *pHML-P2A-Gal4* plasmid (gift from A. Herzig^27^) was PCR-amplified and cloned into *pattB-LHG* (gift from L. Mathies). The transgene was inserted into the *su(Hw)attP8* site on the X chromosome. To build *LexAop-NLSmScarlet3*, codon-optimized *3xNLS-mScarlet3* cDNA was synthesized by Integrated DNA Technologies (IDT) and cloned into *pJFRC19*. The transgene was inserted into the *su(Hw)attP5* site on chromosome II.

## METHOD DETAILS

### Ovarian tumor induction, lifespan assays, and tumor progression

Upon eclosion, adult females were placed at 21°C on food with yeast powder for two days. A maximum of 25 flies were kept in each vial. Flies were then moved to new food and shifted to 29°C to initiate tumor induction. Flies were transferred to new food every two days, and the number of dead flies was counted. For each lifespan assay, at least 50 flies were used for each sample group, with control and experimental groups assessed contemporaneously. OC tumors were graded as described previously^25^ on a scale of 1 to 4 with increasing severity.

### Immunofluorescence

Tissues were dissected in PBS and fixed in 4% PFA-PBS for one hour at room temperature without agitation. Samples were washed three times with PBS-TX (0.1% Triton-X in 1X PBS). For primary antibody staining, samples were blocked in 2% BSA for 30-60 minutes. Samples were incubated overnight in primary antibody at 4C at the following concentrations: NimC1(1:100), Dcp1 (1:100), MMP1 (1:100), Laminin (1:500), Flag M2 (1:100). Samples were washed three times with PBS-TX and incubated with AlexaFluor-conjugated secondary antibodies for 90 minutes at room temperature. To stain actin, fixed tissues were incubated for 60-90 minutes at room temperature with Rhodamine-Phalloidin at a concentration of 1:500. To stain nuclei, a 1:5000 DAPI solution was applied for 15 minutes. After all incubations, samples were washed three times with PBS. Samples were mounted in Diamond Antifade Mountant before imaging. For ROS visualization, DHE staining was performed as described previously^37^. Dissected unfixed tissues were incubated in 30 uM DHE solution for 5 min, washed three times with PBS followed by mounting and imaging immediately.

### Protease activity assays

Tissues were rapidly dissected in Schneider’s media and transferred into a solution of 20% casein or gelatin in media. Tubes were gently agitated for 25 minutes, washed with PBS, and fixed in 4% PFA. Following further washing, samples were stained with primary antibody (if necessary) or only phalloidin and DAPI. Samples were imaged within 72 hours to avoid degradation.

### Confocal microscopy and image analysis

Fixed samples were imaged on the Zeiss LSM900 Confocal Microscope with a Plan-APOCHROMAT 20x/0,8 objective. Images were processed and analyzed with FIJI software. Unless otherwise indicated, all images are maximum projections. Maximum projection images were created in FIJI software.

### Statistics

The one-way ANOVA test or Student’s unpaired T-test were used for parametric data. The Mann-Whitney U test was used for tumor progression. The Log-Rank test was used to determine significant differences in survival curves. Graphpad Prism was used to perform statistical testing. All error bars show S.E.M.

### Disc Wounds

For ex vivo wounding, wing discs were dissected from 3rd instar larvae in culture medium (15% FBS, 10,000 u/mL penicillin/streptomycin in Schneider’s medium), physically pinched by a sharp forceps, and incubated at 29°C for three hours before fixation or protease activity assays. For in vivo wounding, 3rd instar larval wing discs expressing *en-Gal4>UAS-RFP* were forceps-pinched in cold PBS under an RFP dissecting scope. After pinching, larvae were transferred to a food vial and kept at 29°C for three hours before dissection.

### Macrophage Isolation

To isolate circulating macrophages, individual female flies expressing *Hml*^*P2A*^*-LHG>LexAop-NLSmScarlet3* were bled by opening the abdomen in a PBS droplet on a poly-D-lysine coated glass bottom dish. Cells were allowed to settle for 15 minutes and fixed in 4% PFA for 15 minutes. The carcass was then separated into head, thorax/legs, and abdomen to isolate sessile populations. Ovaries were removed from the abdomen and stored separately. All tissues were separately fixed in 4% PFA for 15 minutes.

### Macrophage signal measurement

The NimC1 signal on ovaries (excluding vitellogenic follicles) or wing discs was measured using FIJI. A threshold was set using FIJI’s *Threshold* function to remove background signal, and the resulting area of the NimC1 signal was quantified. The relative area was then calculated by dividing each value by the mean value of the control RNAi group.

## FIGURE LEGENDS

**Supplementary Figure 1: (A)** Tumor-specific knockdown of *Tep3* with either of two RNAi constructs extends host lifespan. **(B-C)** Alphafold rendering of Tep3 **(B)** and its mammalian homolog CD109 **(C)** highlights structural similarities between the proteins. (D) *Tep3* KD in WT ovaries does not affect macrophage association (n-values: Ovary =12, Ovary + *Tep3* KD = 12). Log-rank test (A) and unpaired t-test (D) used to determine significance; *p < .05, ***p < .0005. Dots in (D) represent individual samples.

**Supplementary Figure 2: (A, B)** At day 20 ATI, *Tep3* KD does not change the amount of apoptosis in the tumor, indicated by Dcp1 staining (pink), quantitated in **(C)** (n values: Control = 11, Control + *Tep3* KD = 21, Tumor = 29, Tumor + *Tep3* KD = 16). **(D)** Inducing apoptosis in follicle cells increases apoptosis but is not sufficient to recruit macrophages **(E). (F-H)** Similarly, inhibiting apoptosis in tumors does not affect macrophage recruitment, quantitated in **(I)** (n values: Tumor = 9, Tumor + *p35* OE = 9, Tumor + *Diap1* OE = 14). **(K-L)** Expressing *Duox* RNAi in tumors decreases ROS levels but does not change macrophage association **(K’-L’)**, assayed by *pxn-LexA>LexAop-mCherry*), quantitated in **M, N** (n values: Tumor = 10, Tumor + *Duox* KD = 10). Scale bars = 50μm; Unpaired t-test (M, N) and One Way ANOVA (C, I) used to determine significance; ****p < .00005. Dots in (C, I, M, N) represent individual samples.

**Supplementary Figure 3:** WT ovaries with *Col4* KD **(A, B)** or *lanB* KD **(D, E)** show slight increase in macrophage recruitment (green), quantitated in **C** (n values: Ovary = 23, Ovary + *Col4* KD = 18) and **F** (n values: Ovary = 33, Ovary + *lanB* KD = 25). Scale bars = 50μm; Unpaired t-test (C, F) used to determine significance; *p < .05, **p < .005. Dots in (C, F) represent individual samples. **(G, H)** Tumors display increased Casein-FITC fluorescence (green) compared to WT ovaries. Tumor Casein-FITC signal (green) **(I)** is correlated with MMP1 expression (anti-MMP1 antibody, white) **(I’)**. Scale bars = 50μm; Unpaired t-test (C, F) used to determine significance. *p < .05, **p < .005. Dots in (C, F) represent individual samples.

**Supplementary Figure 4: (A, B)** *Tep3* KD in tumors does not change MMP2 activity, indicated by Gelatin-FITC fluorescence (white), quantitated in **C** (Tumor = 27, Tumor + *Tep3* KD = 39). **(D)** Complementation testing of *Tep3* alleles indicates that *TepqΔ* allele is hypomorphic for *Tep3* function. Scale bars = 50μm; Unpaired t-test (C) used to determine significance. Dots in (C) represent individual samples.

**Supplementary Table S1:** Experimental genotypes

