## Supplemental Figures for "Fly wounds and tumors restrict macrophages via a matrix degradation-moderating protease inhibitor"

**FIGURE S1:**

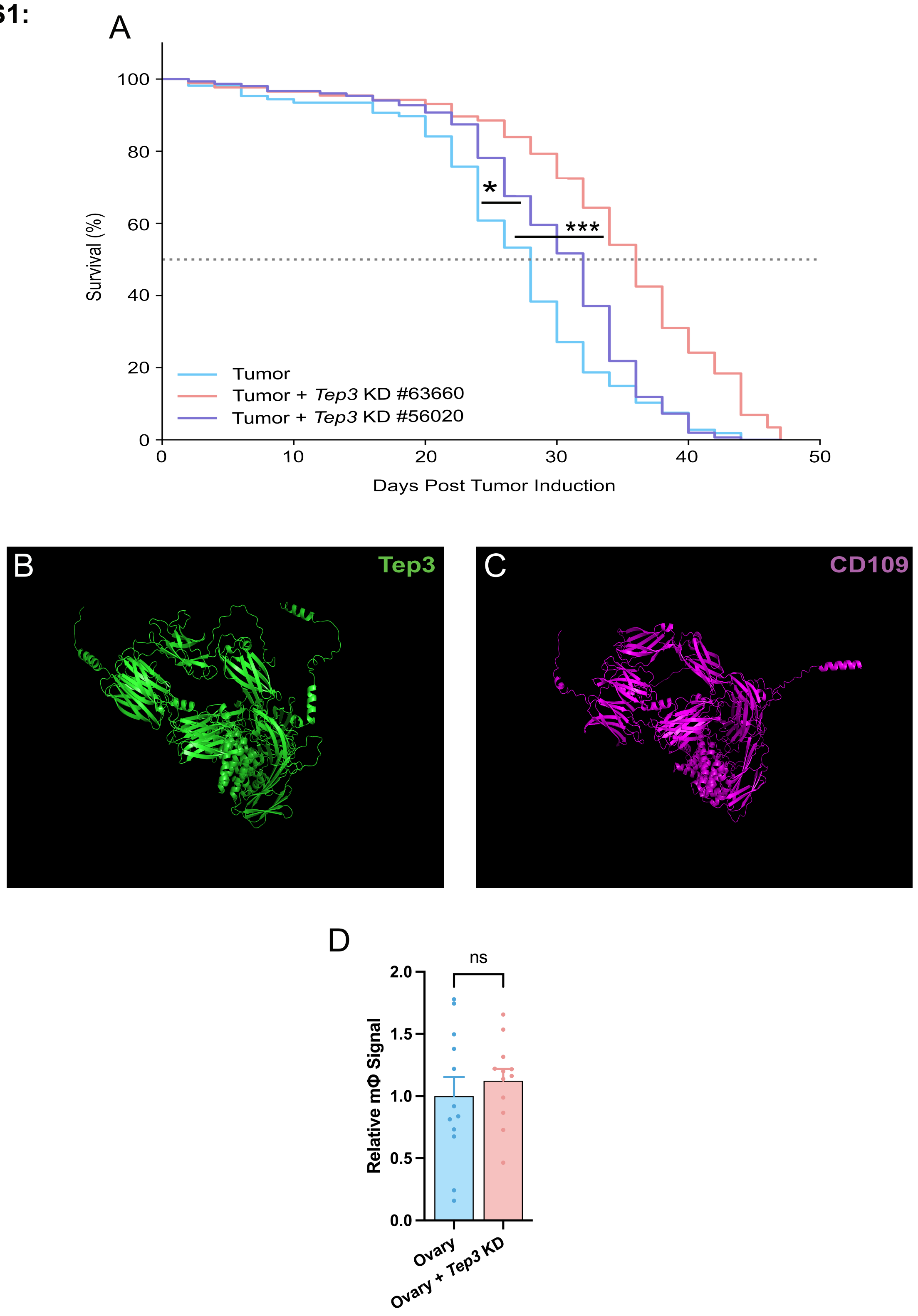

A
100
80
60
40
20
0
Survival (%)
*
***
Tumor
Tumor + Tep3 KD #63660
Tumor + Tep3 KD #56020
0
10
20
30
40
50
Days Post Tumor Induction
B
Tep3
C
CD109
D
ns
2.0
1.5
1.0
0.5
0.0
Relative mΦ Signal
Ovary
Ovary + Tep3 KD

**FIGURE S2:**

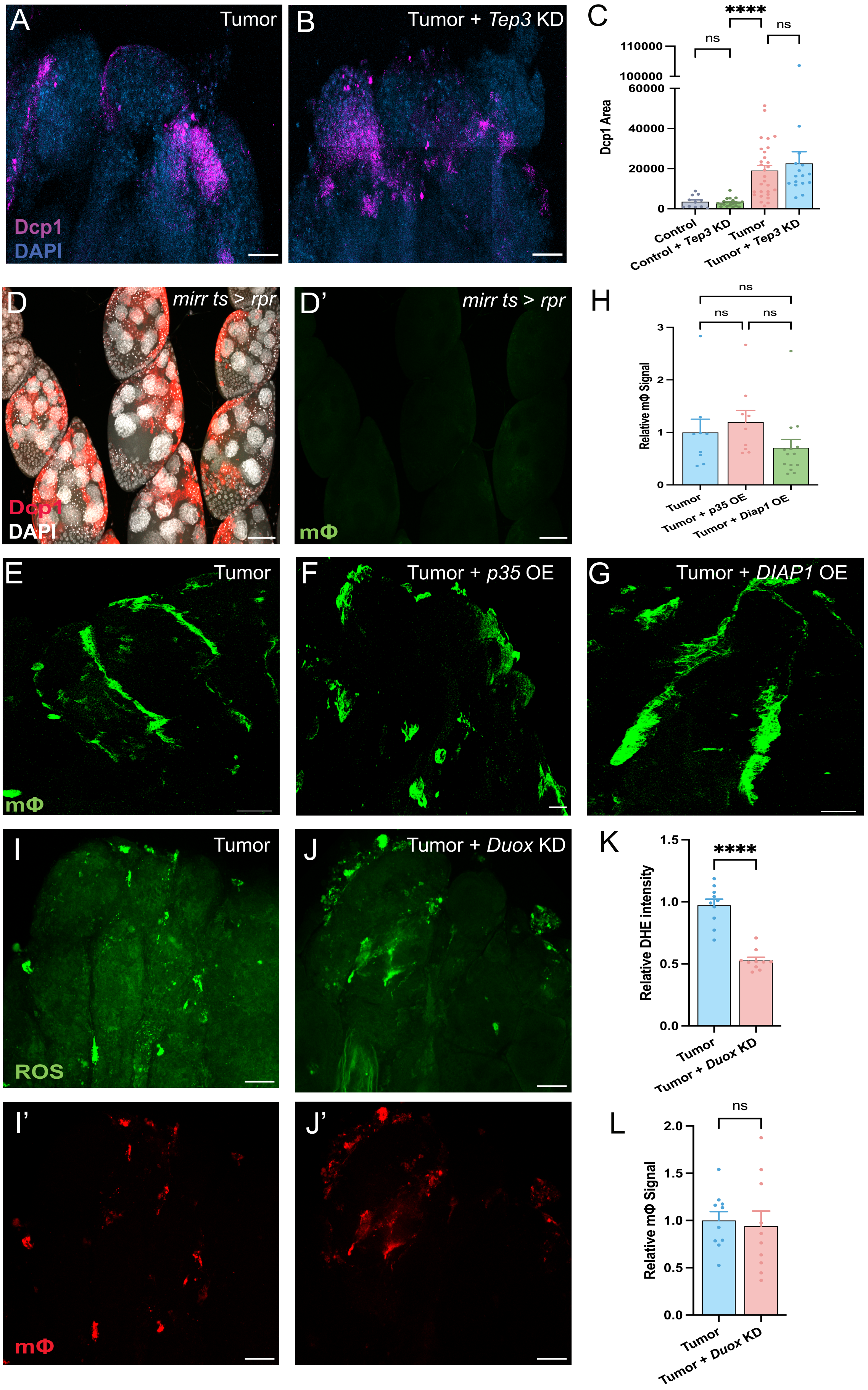

A
Tumor
Dcp1
DAPI
B
Tumor + Tep3 KD
C
****
ns
ns
110000
100000
60000
40000
20000
0
Dcp1 Area
Control
Control + Tep3 KD
Tumor
Tumor + Tep3 KD
D
mirr ts > rpr
Dcp1
DAPI
D'
mirr ts > rpr
mΦ
H
ns
ns
ns
3
2
1
0
Relative mΦ Signal
Tumor
Tumor + p35 OE
Tumor + Diap1 OE
E
Tumor
mΦ
F
Tumor + p35 OE
G
Tumor + DIAP1 OE
I
Tumor
ROS
J
Tumor + Duox KD
K
****
1.5
1.0
0.5
0.0
Relative DHE intensity
Tumor
Tumor + Duox KD
I'
mΦ
J'
L
ns
2.0
1.5
1.0
0.5
0.0
Relative mΦ Signal
Tumor
Tumor + Duox KD

**FIGURE S3:**

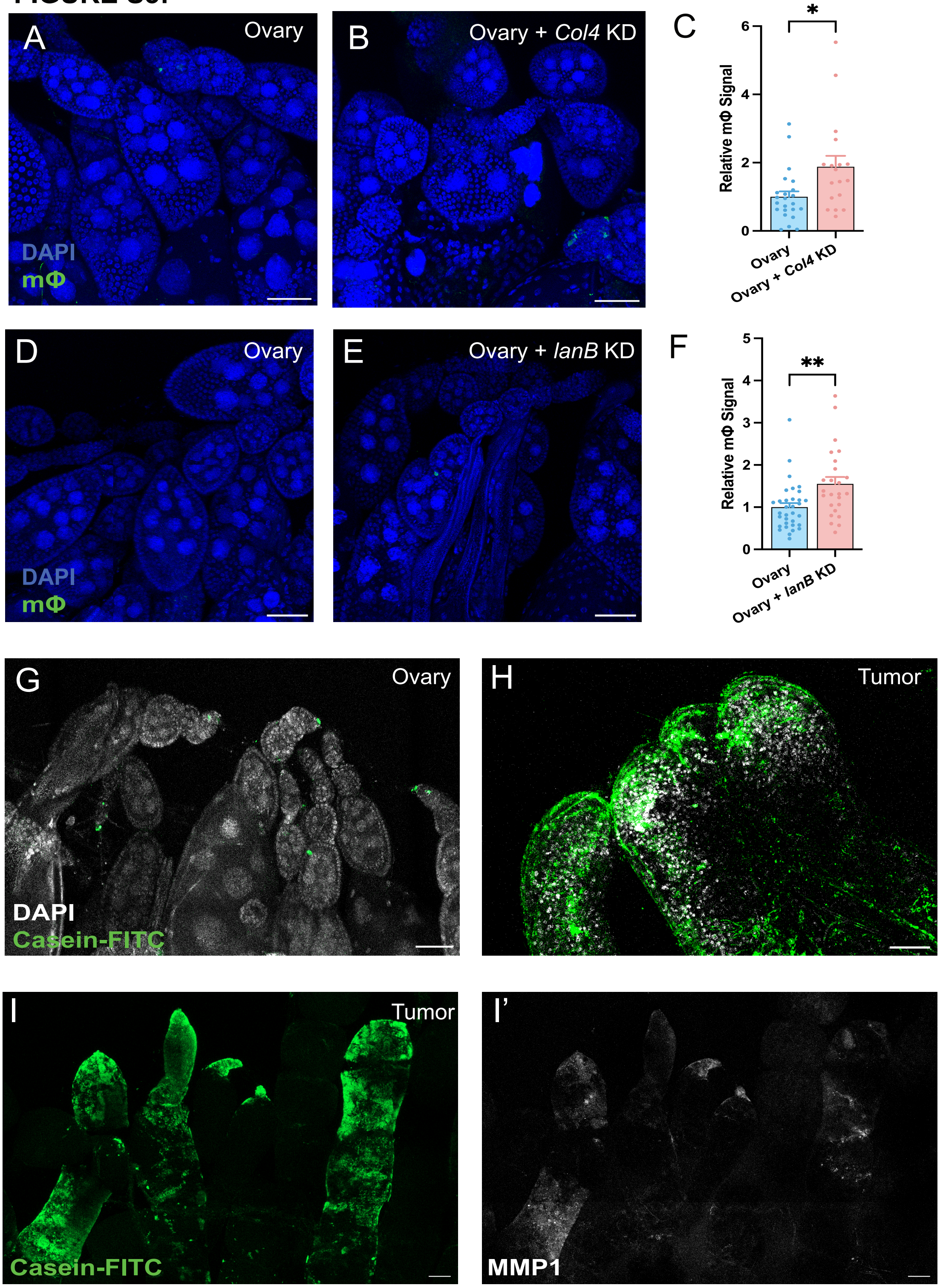
A
Ovary
DAPI
mΦ
B
Ovary + Col4 KD
C
Relative mΦ Signal
*
0
2
4
6
Ovary
Ovary + Col4 KD
D
Ovary
DAPI
mΦ
E
Ovary + lanB KD
F
Relative mΦ Signal
**
0
1
2
3
4
5
Ovary
Ovary + lanB KD
G
Ovary
DAPI
Casein-FITC
H
Tumor
I
Tumor
Casein-FITC
I'
MMP1

FIGURE S4:

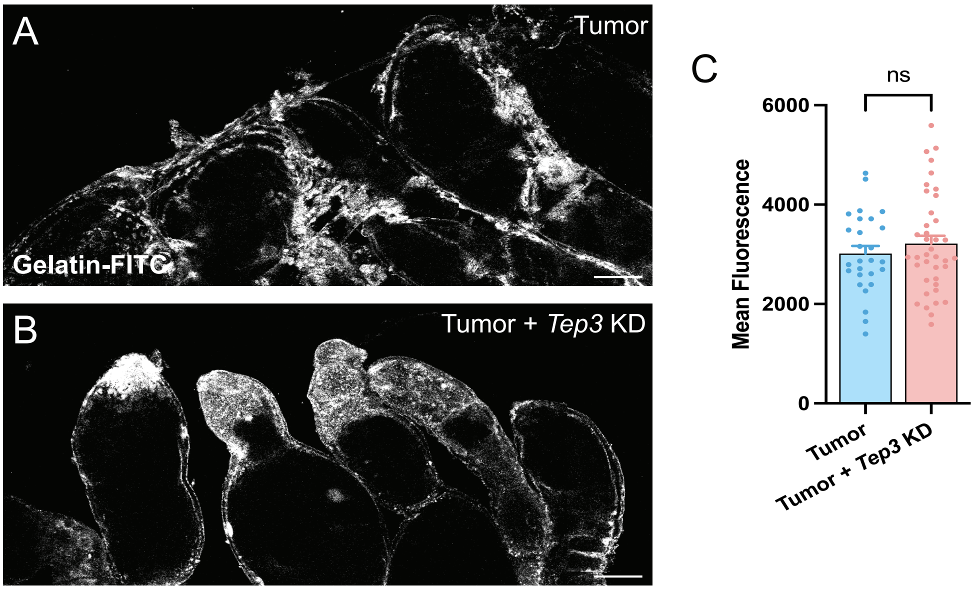


D

| | ***Tep3$^{sk2}$*** | ***Tep3 GAL4*** | ***TepqΔ*** |
|---|---|---|---|
| ***Tep3$^{sk2}$*** | Lethal | Lethal | Viable |
| ***Tep3 GAL4*** | Lethal | Lethal | Viable |
| ***TepqΔ*** | Viable | Viable | Viable |

**TABLE S1:**

| Figure | Panel | Condition | Genotype |
|---|---|---|---|
| **1** | A, B, I | Control | *Hml$^{P2A}$-LHG/+; tj-Gal4, tubulin-Gal80$^{ts}$/LexAop-3xnls::mScarlet3; +/+* |
| | | Tumor | *Hml$^{P2A}$-LHG/+; tj-Gal4, tubulin-Gal80$^{ts}$/LexAop-3xnls::mScarlet3; UAS-aPKC$^{\Delta N}$, UAS-Ras$^{V12}$/+* |
| | D-G | Tumor | *tj-Gal4, tubulin-Gal80$^{ts}$/+; UAS-aPKC$^{\Delta N}$, UAS-Ras$^{V12}$/+* |
| | J, K, M | Tumor | *tj-LHG, tubulin-Gal80$^{ts}$/+; LexAop-aPKC$^{\Delta N}$-T2A-Ras$^{V12}$, Hml$^{P2A}$-Gal4/+* |
| | | Tumor + no macrophages | *tj-LHG, tubulin-Gal80$^{ts}$/UAS-hid:UAS-rpr; LexAop-aPKC$^{\Delta N}$-T2A-Ras$^{V12}$, Hml$^{P2A}$-Gal4/+* |
| **2** | A-C | WT | *tubulin-Gal80$^{ts}$/Tep3-Gal4; +/UAS-mCD8GFP* |
| | | Tumor | *tj-LHG, tubulin-Gal80$^{ts}$/Tep3-Gal4; LexAop-aPKC$^{\Delta N}$-T2A-Ras$^{V12}$/UAS-mCD8GFP* |
| | E-G, I, J | Control | *tj-Gal4, tubulin-Gal80$^{ts}$/UAS-GFP KD; +/+* |
| | | Tumor | *tj-Gal4, tubulin-Gal80$^{ts}$/UAS-GFP KD; UAS-aPKC$^{\Delta N}$, UAS-Ras$^{V12}$/+* |
| | | Tumor + Tep3 KD | *tj-Gal4, tubulin-Gal80$^{ts}$/UAS-Tep3 KD; UAS-aPKC$^{\Delta N}$, UAS-Ras$^{V12}$/+* |
| | L | en>RFP + Tep3 OE | *en-Gal4, UAS-RFP/+; Dr/UAS-Tep3-FLAG* |
| | M | Tumor | *tj-Gal4, tubulin-Gal80$^{ts}$/UAS-GFP KD; UAS-aPKC$^{\Delta N}$, UAS-Ras$^{V12}$/LexAop-rpr* |
| | | Tumor + Tep3 KD | *tj-Gal4, tubulin-Gal80$^{ts}$/UAS-Tep3 KD; UAS-aPKC$^{\Delta N}$, UAS-Ras$^{V12}$/LexAop-rpr* |
| | | Tumor (mφ depleted) | *Hml$^{P2A}$-LHG/+; tj-Gal4, tubulin-Gal80$^{ts}$/UAS-GFP KD; UAS-aPKC$^{\Delta N}$, UAS-Ras$^{V12}$/LexAop-rpr* |
| | | Tumor + Tep3 KD (mφ depleted) | *Hml$^{P2A}$-LHG/+; tj-Gal4, tubulin-Gal80$^{ts}$/UAS-Tep3 KD; UAS-aPKC$^{\Delta N}$, UAS-Ras$^{V12}$/LexAop-rpr* |
| **3** | A, B | Tumor | *tj-Gal4, tubulin-Gal80$^{ts}$/UAS-GFP KD; UAS-aPKC$^{\Delta N}$, UAS-Ras$^{V12}$/+* |
| | | Tumor + Tep3 KD | *tj-Gal4, tubulin-Gal80$^{ts}$/UAS-Tep3 KD; UAS-aPKC$^{\Delta N}$, UAS-Ras$^{V12}$/+* |
| | D, E | Control | *tj-Gal4, tubulin-Gal80$^{ts}$/vkg-GFP; +/+* |
| | | TJ>MMP1 OE | *tj-Gal4, tubulin-Gal80$^{ts}$/vkg-GFP; UAS-MMP1/+* |
| | H, I | Tumor | *tj-Gal4, tubulin-Gal80$^{ts}$/UAS-GFP KD; UAS-aPKC$^{\Delta N}$, UAS-Ras$^{V12}$/+* |
| | | Tumor + Col4 KD | *tj-Gal4, tubulin-Gal80$^{ts}$/UAS-vkg KD; UAS-aPKC$^{\Delta N}$, UAS-Ras$^{V12}$/UAS-cg25c KD* |
| | N, O | Tumor | *tj-Gal4, tubulin-Gal80$^{ts}$/UAS-GFP KD; UAS-aPKC$^{\Delta N}$, UAS-Ras$^{V12}$/+* |
| | | Tumor + lanB KD | *tj-Gal4, tubulin-Gal80$^{ts}$/UAS-lanB1 KD; UAS-aPKC$^{\Delta N}$, UAS-Ras$^{V12}$/UAS-lanB2 KD* |
| **4** | A-C | Tumor | *tj-Gal4, tubulin-Gal80$^{ts}$/UAS-GFP KD; UAS-aPKC$^{\Delta N}$, UAS-Ras$^{V12}$/+* |
| | | Tumor + MMP1 KD | *tj-Gal4, tubulin-Gal80$^{ts}$/UAS-GFP KD; UAS-aPKC$^{\Delta N}$, UAS-Ras$^{V12}$/UAS-MMP1 KD* |
| | | Tumor + Tep3 KD | *tj-Gal4, tubulin-Gal80$^{ts}$/UAS-Tep3 KD; UAS-aPKC$^{\Delta N}$, UAS-Ras$^{V12}$/+* |
| | E, F | Tumor | *tj-Gal4, tubulin-Gal80$^{ts}$/UAS-GFP KD; UAS-aPKC$^{\Delta N}$, UAS-Ras$^{V12}$/+* |
| | | Tumor + Tep3 KD | *tj-Gal4, tubulin-Gal80$^{ts}$/UAS-Tep3 KD; UAS-aPKC$^{\Delta N}$, UAS-Ras$^{V12}$/+* |
| | H-K | Tumor | *tj-Gal4, tubulin-Gal80$^{ts}$/UAS-GFP KD; UAS-aPKC$^{\Delta N}$, UAS-Ras$^{V12}$/+* |
| | | Tumor + MMP1 KD | *tj-Gal4, tubulin-Gal80$^{ts}$/UAS-GFP KD; UAS-aPKC$^{\Delta N}$, UAS-Ras$^{V12}$/UAS-MMP1 KD* |
| | | Tumor + Tep3 KD | *tj-Gal4, tubulin-Gal80$^{ts}$/UAS-Tep3 KD; UAS-aPKC$^{\Delta N}$, UAS-Ras$^{V12}$/+* |
| | | Tumor + MMP1 KD + Tep3 KD | *tj-Gal4, tubulin-Gal80$^{ts}$/UAS-Tep3 KD; UAS-aPKC$^{\Delta N}$, UAS-Ras$^{V12}$/UAS-MMP1 KD* |
| | N, O | Tumor | *tj-Gal4, tubulin-Gal80$^{ts}$/UAS-GFP KD; UAS-aPKC$^{\Delta N}$, UAS-Ras$^{V12}$/+* |
| | | Tumor + Tep3 KD | *tj-Gal4, tubulin-Gal80$^{ts}$/UAS-Tep3 KD; UAS-aPKC$^{\Delta N}$, UAS-Ras$^{V12}$/+* |

| | | | |
|---|---|---|---|
| | Q, R | Control | *tubulin-Gal80$^{ts}$/+; Hml$^{P2A}$-Gal4/UAS-mCherry KD* |
| | | Tep3 OE | *tubulin-Gal80$^{ts}$/+; Hml$^{P2A}$-Gal4/UAS-Tep3-FLAG* |
| **5** | A, B | Unwounded/Wounded | *Tep3-Gal4/+; UAS-mCD8GFP/+* |
| | C, D | Unwounded/Wounded | *en-Gal4, UAS-RFP/+; Dr/+* |
| | F, H, J | en>RFP + Tep3 OE | *en-Gal4, UAS-RFP/+; Dr/UAS-Tep3-FLAG* |
| **S1** | A | Tumor | *tj-Gal4, tubulin-Gal80$^{ts}$/UAS-GFP KD; UAS-aPKC$^{\Delta N}$, UAS-Ras$^{V12}$/+* |
| | | Tumor + Tep3 KD #63660 | *tj-Gal4, tubulin-Gal80$^{ts}$/UAS-Tep3 KD; UAS-aPKC$^{\Delta N}$, UAS-Ras$^{V12}$/+* |
| | | Tumor + Tep3 KD #56020 | *tj-Gal4, tubulin-Gal80$^{ts}$/UAS-Tep3 KD (#56020); UAS-aPKC$^{\Delta N}$, UAS-Ras$^{V12}$/+* |
| | D | Ovary | *tj-Gal4, tubulin-Gal80$^{ts}$/UAS-GFP KD; +/+* |
| | | Ovary + Tep3 KD | *tj-Gal4, tubulin-Gal80$^{ts}$/UAS-Tep3 KD; +/+* |
| **S2** | A-C | Control | *tj-Gal4, tubulin-Gal80$^{ts}$/UAS-GFP KD; +/+* |
| | | Control + Tep3 KD | *tj-Gal4, tubulin-Gal80$^{ts}$/UAS-Tep3 KD; +/+* |
| | | Tumor | *tj-Gal4, tubulin-Gal80$^{ts}$/UAS-GFP KD; UAS-aPKC$^{\Delta N}$, UAS-Ras$^{V12}$/+* |
| | | Tumor + Tep3 KD | *tj-Gal4, tubulin-Gal80$^{ts}$/UAS-Tep3 KD; UAS-aPKC$^{\Delta N}$, UAS-Ras$^{V12}$/+* |
| | D | mirr ts > rpr | *UAS-rpr, tubulin-Gal80$^{ts}$/+; mirr-Gal4/+* |
| | F-H | Tumor | *tj-Gal4, tubulin-Gal80$^{ts}$/UAS-luciferase; UAS-aPKC$^{\Delta N}$, UAS-Ras$^{V12}$/+* |
| | | Tumor + p35 OE | *tj-Gal4, tubulin-Gal80$^{ts}$/Sp; UAS-aPKC$^{\Delta N}$, UAS-Ras$^{V12}$/UAS-p35* |
| | | Tumor + DIAP1 OE | *tj-Gal4, tubulin-Gal80$^{ts}$/+; UAS-aPKC$^{\Delta N}$, UAS-Ras$^{V12}$/UAS-DIAP1* |
| | K, L | Tumor | *tj-Gal4, tubulin-Gal80$^{ts}$/Pxn-LexA, LexAop-mCherry; UAS-aPKC$^{\Delta N}$, UAS-Ras$^{V12}$/UAS-GFP KD* |
| | | Tumor + Duox KD | *tj-Gal4, tubulin-Gal80$^{ts}$/Pxn-LexA, LexAop-mCherry; UAS-aPKC$^{\Delta N}$, UAS-Ras$^{V12}$/UAS-Duox KD* |
| **S3** | A, B | Ovary | *tj-Gal4, tubulin-Gal80$^{ts}$/UAS-GFP KD; +/+* |
| | | Ovary + Col4 KD | *tj-Gal4, tubulin-Gal80$^{ts}$/UAS-vkg KD; +/UAS-cg25c KD* |
| | D, E | Ovary | *tj-Gal4, tubulin-Gal80$^{ts}$/UAS-GFP KD; +/+* |
| | | Ovary + lanB KD | *tj-Gal4, tubulin-Gal80$^{ts}$/UAS-lanB1 KD; +/UAS-lanB2 KD* |
| | G, H | Ovary | *tj-Gal4, tubulin-Gal80$^{ts}$/w1118; +/+* |
| | | Tumor | *tj-Gal4, tubulin-Gal80$^{ts}$/UAS-GFP KD; UAS-aPKC$^{\Delta N}$, UAS-Ras$^{V12}$/+* |
| | I, I' | Tumor | *tj-Gal4, tubulin-Gal80$^{ts}$/UAS-GFP KD; UAS-aPKC$^{\Delta N}$, UAS-Ras$^{V12}$/+* |
| **S4** | A, B | Tumor | *tj-Gal4, tubulin-Gal80$^{ts}$/UAS-GFP KD; UAS-aPKC$^{\Delta N}$, UAS-Ras$^{V12}$/+* |
| | | Tumor + Tep3 KD | *tj-Gal4, tubulin-Gal80$^{ts}$/UAS-Tep3 KD; UAS-aPKC$^{\Delta N}$, UAS-Ras$^{V12}$/+* |
